# High Resolution Imaging Mass Spectrometry of Bacterial Microcolonies at Ecological Scales

**DOI:** 10.1101/717066

**Authors:** Rita de Cassia Pessotti, Bridget L. Hansen, Vineetha M. Zacharia, Daniel Polyakov, Matthew F. Traxler

**Author notes:** Correspondence, University of California, Berkeley, 111 Koshland Hall, Berkeley CA 94720.

## Abstract

Microbes interact with the world around them at the chemical level. However, directly examining the chemical exchange between microbes, and microbes and their environment, at ecological scales, *i.e.* the scale of a single bacterial cell or small groups of cells, remains a key challenge. Here we address this obstacle by presenting a methodology that enables <u>M</u>atrix-<u>a</u>ssisted <u>l</u>aser <u>d</u>esorption/<u>i</u>onization (MALDI) <u>i</u>maging <u>m</u>ass <u>s</u>pectrometry (IMS) of bacterial microcolonies. By combining optimized sample preparation with sub-atmospheric pressure MALDI, we demonstrate that chemical output from groups of as few as ~50 cells can be visualized with MALDI-IMS. Application of this methodology to *Bacillus subtilis* and *Streptomyces coelicolor* revealed heterogeneity in chemical output across microcolonies, and asymmetrical metabolite production when cells grew within physiological gradients produced by *Medicago sativa* roots. Taken together, these results indicate that MALDI-IMS can readily visualize metabolites made by very small assemblages of bacterial cells, and that even these small groups of cells can differentially produce metabolites in response to local chemical gradients.

## Body

In microbial habitats, from soils to oceans to the intestinal lumen, nutrient availability and physical topology are heterogenous, and chemical gradients are pervasive (1–4). Our knowledge of the spatial distribution of bacteria within these environments is relatively sparse, however, it is widely hypothesized that bacteria exist as single cells or in small patches of clonal populations (1, 4-6). Thus, microbes likely interact with environmental gradients and each other at scales on the order of microns to hundreds of microns.

Microbes may modify their local chemical environments in multiple ways, including depleting nutrients, excreting metabolic waste, and producing public goods such as siderophores and biofilm matrices (7–9). Many microbes also make a wide array of specialized metabolites, which are thought to mediate interactions ranging from chemical warfare to cooperative coordination (10, 11). Additionally, bacteria may modify the lipid content of their membranes depending on environmental conditions (12–14). These molecules likely function in a world of microgradients, though what this may look like in space remains largely unknown. Gaining an understanding of how microbes respond to, and alter, their local chemical environments, including within microbiomes, will require methodologies that provide spatial information about microbial chemistry at micron scales.

<u>M</u>atrix-<u>a</u>ssisted <u>l</u>aser <u>d</u>esorption/<u>i</u>onization (MALDI) <u>i</u>maging <u>m</u>ass <u>s</u>pectrometry (IMS) is a promising technique for assessing spatial patterns of microbial chemistry (15–17). To this point, direct MALDI-IMS of microbes has mostly been applied to macroscale colonies. These structures are relatively large (e.g. millimeters in diameter), and spatial resolutions of ~400-600 µm pixels have been sufficient to visualize differential metabolite production (15–17). However, the limits of MALDI-IMS remain to be explored in terms of visualizing molecules produced by bacteria at ecologically-relevant spatial scales, i.e. on the order of 1-100 µm. Here we report our progress in using IMS to visualize metabolites produced by microcolonies of bacteria, at a high spatial resolution of 10 µm.

To determine if features from microcolonies of bacteria could be detected by MALDI-IMS, we inoculated the well-characterized organisms *Bacillus subtilis* 3610 and *Streptomyces coelicolor* M145 directly onto thin pads of agarose mounted on <u>i</u>ndium-<u>t</u>in <u>o</u>xide (ITO) coated microscope slides. These microbes were selected because they produce several well-characterized families of specialized metabolites and membrane lipids under standard laboratory conditions. Strains were inoculated onto agarose pads in 15 µL volumes of 25% strength ISP2 medium and incubated for 14 h (*B. subtilis*) or 48 h (*S. coelicolor*). This allowed growth of individual *B. subtilis* cells into microcolonies of cell chains in the range of 100-200 µm long, and microcolonies of *S. coelicolor* in the range of 100-300 µm in diameter. Micrographs of the colonies were acquired using a phase contrast microscope, the samples were dried, and MALDI matrix was sublimated onto the surface of the sample to achieve a uniform layer of fine matrix crystals. The sample was then imaged with a pixel size of 10 µm using a sub-atmospheric pressure MALDI source equipped with an ion funnel (18) coupled to an Orbitrap Q-Exactive mass spectrometer, a combination that provides both high spatial and high mass resolution.

Analysis of the MALDI images revealed a variety of chemical features with spatial distributions that matched the cellular arrangement seen in the phase contrast micrographs with surprising fidelity (Fig. 1A and B, **supplementary Fig. S2A-K, S5A-H**). We were able to chemically identify a subset of the features (**Supplementary Tables S1 and S2**) based on exact mass and MS/MS fragmentation patterns. For *B. subtilis* this included the lipopeptide specialized metabolite C15-surfactin [M+H]^+^ (*m/*z 1036.7 ± 0.1) (**Supplementary Fig. S1A-C**) and multiple species of membrane lipids, exemplified by the phosphotidylglycerol (PG) (32:0) [M+Na]+ (*m/*z 745.5 ± 0.1) (**Supplementary Fig. S1E, F and Fig. S3**). The MS/MS fragmentation pattern of the ion with the *m/*z 1036.7 ± 0.1 detected in a sample of *B. subtilis* macrocolony was identical to that of an authentic commercial standard (Fig. 1B). Under these conditions, surfactin was only detectable in the largest microcolonies, while membrane lipids were detectable in almost all microcolonies (Fig. 1A, **supplementary Fig. S2A-K**). Given the known size range for *B. subtilis* cells (19), we calculate that the colony marked with a black arrow in Fig. 1A is comprised of 26-52 cells.

**Fig. 1:**
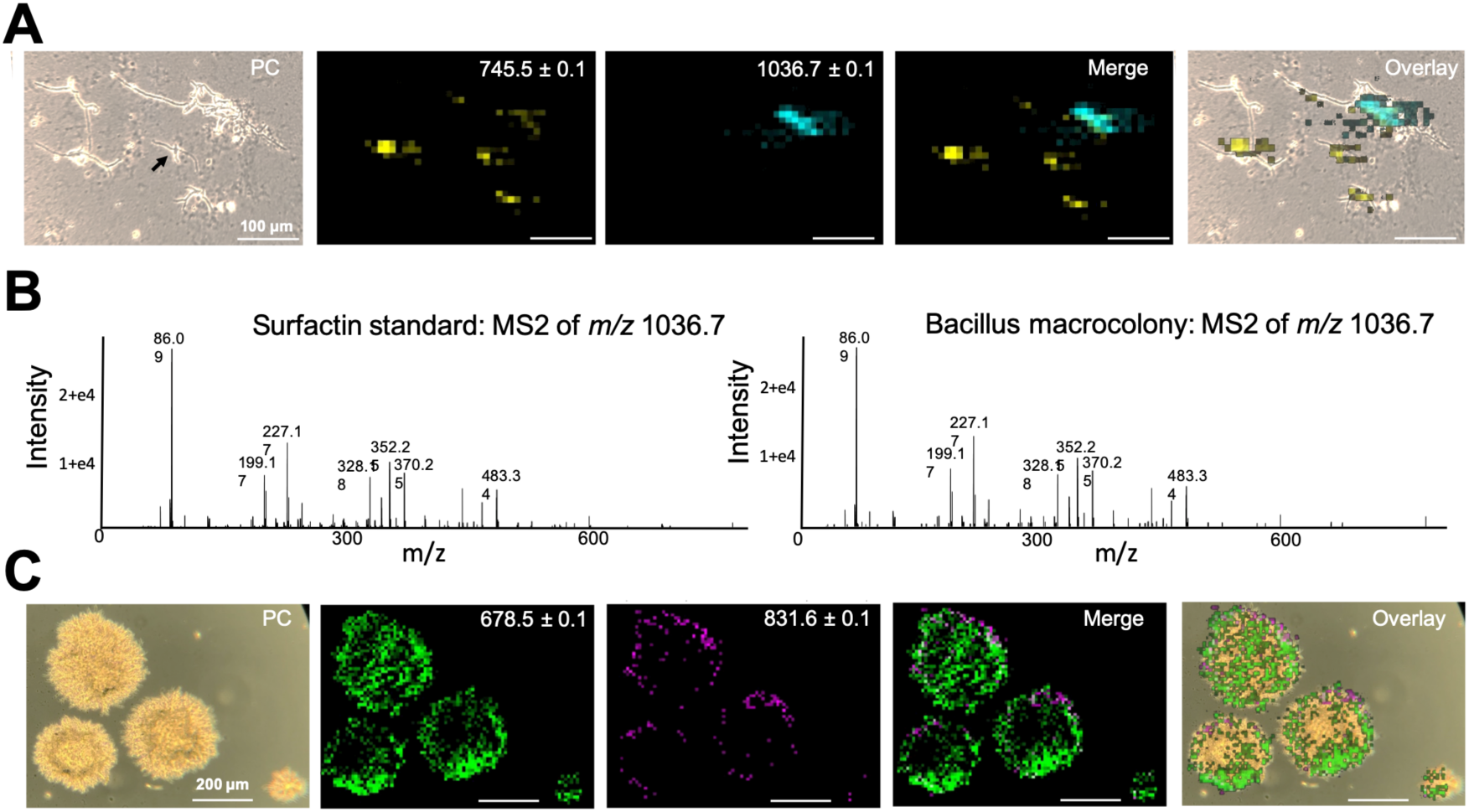
Features associated with bacterial microcolonies are detectable at micron scales by MALDI-IMS. **A)** Micrograph of *B. subtilis* microcolonies and MALDI images showing the spatial distribution of two of the detected features: *m/z* 745.5 ± 0.1 [sodium adduct of the lipid PG (32:0)] and *m/z* 1036.7 ± 0.1 (proton adduct of C15-surfactin). Scale bar = 100 microns. **B)** MS2 spectra acquired for the ion with the *m/z* 1036.7 from a commercial surfactin standard and *B. subtilis* macrocolony showing identical fragmentation patterns. **C)** Micrograph of *S. coelicolor* microcolonies and MALDI images showing the spatial distribution of two detected features: *m/z* 678.4 ± 0.1 [putative proton adduct of the lipid PE(31:0)] and *m/z* 831.6.4 ± 0.1 (a putative uncharacterized lipid). Scale bar = 200 microns. All micrographs were acquired using a phase contrast (PC) microscope.

For *S. coelicolor* we detected multiple features that correlate with phosphatidylethanolamine (PE) lipids (e.g. *m/*z 678.4 ± 0.1, **Supplementary Fig. S4A**) (13), and a separate series of features previously hypothesized to represent a set of uncharacterized lipids exemplified by *m/z* 831.6 ± 0.1 (20) (**Supplementary Fig. S4B, C and S6**). Interestingly, the PE lipids were distributed across the entirety of the microcolonies, while the uncharacterized lipids were localized to the outer periphery of the colonies (Fig. 1C, **supplementary Fig. S5**). These observations are consistent with the dynamic lipidome made by this organism (13). Together, these results demonstrate that metabolites produced by bacterial microcolonies comprised of relatively low numbers of cells can be readily detected using MALDI-IMS.

The methodology presented here could provide insight into the chemical responses of microbes to physiological microgradients, such as those that likely occur in natural environments. Therefore, we sought to image microcolonies growing in the context of a naturally-produced nutrient gradient. To do so, we inoculated *B. subtilis* and *S. coelicolor* onto slide-mounted agarose pads and laid living *Medicago sativa* roots across them. This setup led to bacterial microcolonies which grew at varying distances from the root, and hence experienced a differential gradient of nutrients available in root exudates.

MALDI-IMS revealed asymmetric production of metabolites across and within bacterial microcolonies within this nutrient gradient. For example, surfactin was produced by microcolonies of *B. subtilis* nearest the root after 14 hours of growth, while membrane lipids were detectable in microcolonies farther away from the root (Fig. 2A, **supplementary Fig. S1D, G and S2L-V**). This finding is consistent with the proposed role of surfactin in initiating biofilm formation on plant roots (21–23), although this has never been observed at the chemical level at the microcolony scale. This observation is also in broad agreement with the findings of Debois and co-workers, who saw surfactin produced by *Bacillus amyloliquefaciens* biofilms on tomato and *Arabidopsis* roots after 24 hours of colonization using IMS at a macro scale. (24). Thus, the methodology presented here can shed light on the spatial distribution of the chemistry involved in the very early stages of root colonization, when metabolite biosynthesis is heterogenous at the level of individual microcolonies. We also found that the different surfactin adducts [M+H]^+^, [M+Na]^+^, and [M+K]^+^ showed subtly distinct distributions (Fig. 2A and **supplementary Fig. S2U and V**), a phenomenon that was also observed by Debois *et. al.* (23, 24) indicating that the root likely generated a microgradient of cations.

**Fig. 2:**
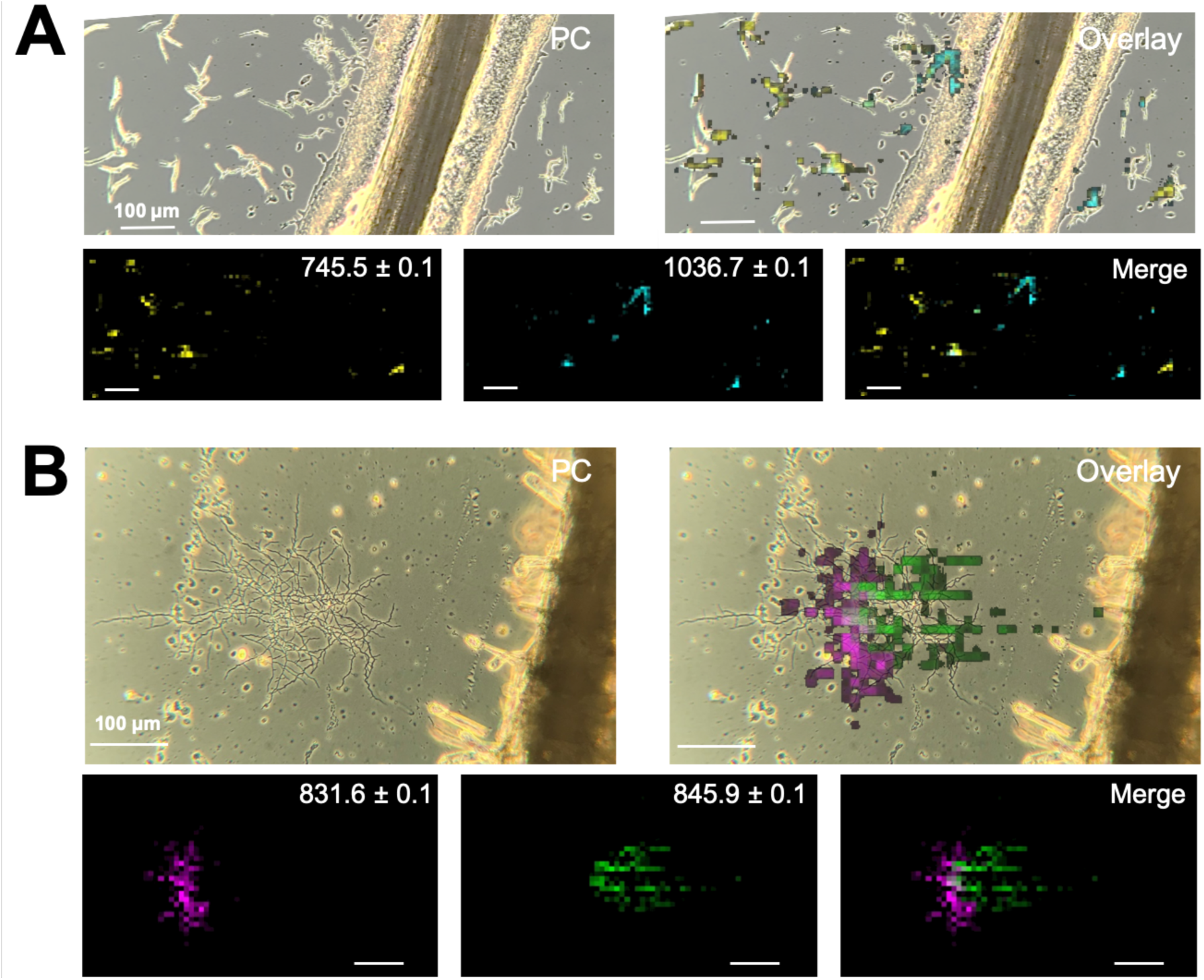
Bacterial microcolonies display asymmetrical metabolite production when grown in proximity to *Medicago sativa* roots. **A)** Micrograph of *B. subtilis* microcolonies growing near a *M. sativa* root; MALDI images show the spatial distribution of two of the detected features: *m/z* 745.5 ± 0.1 [sodium adduct of the lipid PG (32:0)] and *m/z* 1036.7 ± 0.1 (proton adduct of C15-surfactin). Note that surfactin was detected in microcolonies nearest the root. **B)** Micrograph of a *S. coelicolor* microcolony growing near a *M. sativa* root; MALDI images show the spatial distribution of two of the detected features: *m/z* 831.6 ± 0.1 (a putative uncharacterized lipid) and *m/z* 845.9 ± 0.1 (an uncharacterized compound). Note the differential localization of the two features within the microcolony. Micrographs were acquired using a phase contrast (PC) microscope.

The lipidome of *S. coelicolor* is complex and dynamic (13). Interestingly, an asymmetrical distribution of multiple lipid-associated features was observed within a single *S. coelicolor* microcolony grown in proximity to the root (Fig. 2B). An exemplary feature with *m/z* of 831.6 ± 01 was only detected on the side of the colony facing away from the root. Other members of this putative lipid class showed a similar localization (**Supplementary Fig. S5I-N**). This distribution is in contrast to other colony-associated features that were localized to the side of the colony facing the root (e.g. *m/z* 845.9 ± 0.1). *S. coelicolor* membrane lipid composition has been shown to vary depending on nutrient availability, pH, and osmolarity (13). Thus, the spatially variable lipid profile we observe here likely reflects a tailored response to these types of microgradients produced by the root. This example illustrates the utility of high spatial resolution MALDI-IMS for exploring differential production of cellular components within single bacterial microcolonies.

## Conclusion

The results presented here demonstrate the feasibility of using MALDI-IMS for visualizing microbial chemistry at ecologically-relevant spatial scales. This advance reflects improvements in sample preparation and ionization via sub-atmospheric MALDI, which we report in detail in the materials and methods. This methodology enables the visualization of metabolites differentially produced by individual microbial microcolonies composed of as few as ~50 cells. Beyond this, we found that even small assemblages of bacterial cells can produce metabolites asymmetrically in response to environmental gradients. This advance lays the foundation for interrogating the chemical topologies produced by bacteria within diverse microbiomes. Combining this methodology with other imaging techniques that reveal microbial identity (e.g. fluorescent in situ hybridization) or gene expression (e.g. fluorescent protein promoter fusions) will provide a multi-dimensional view of microbial life *in situ* at microscopic scales.

## Acknowledgements

This research was supported by PAPM-EAGER 1650059 from the National Science Foundation to MFT, and startup funds from University of California, Berkeley to MFT.

