## Supplemental Material for "High Resolution Imaging Mass Spectrometry of Bacterial Microcolonies at Ecological Scales"

Correspondence:

### Materials and Methods

#### Strains and growth conditions

*Bacillus subtilis* 3610 was grown in 5 mL ISP2 overnight at 30ºC, 200 rpm. After 16 h of growth, 0.5 mL was inoculated into 9.5 mL of fresh medium, and incubated again at 30ºC, 200 rpm, until reaching $OD_{600}$ of ~0.5 (3-4h). This culture was diluted 1000x in 25% strength ISP2 medium, and 15 µL of this dilution were spread using an inoculating loop on top of an agarose pad made from 200 µL of 1% agarose solution mounted on an ITO-coated microscope slide, and incubated at room temperature (RT) for 14 h. For *Streptomyces coelicolor* M145 experiments, a spore suspension (15 uL of 5 x $10^4$ CFU/mL in sterile water) was spread onto 25% strength ISP2 medium agarose pad prepared in the same way and incubated at 30ºC for 2 days. The small microcolony denoted with a black arrow in Fig. 1A included two chains of cells with a combined length of ~120 µm. Weart and co-workers found that the median cell length of *B. subtilis* ranges between 2.3 and 4.7 µm, depending on nutrient availability [1]. Thus, we calculate that this microcolony likely contained 26-52 cells.

The same growth conditions were used for MALDI-MS/MS experiments, except for the bacterial inoculum: the *B. subtilis* culture was diluted 10x and the *S. coelicolor* spore suspension concentration was 1 x $10^6$ CFU/mL in order to obtain a lawn of microcolonies to increase the number of detected ions.

#### Root-bacteria interaction experiments

*B. subtilis* and *S. coelicolor* were inoculated on agarose pads mounted on ITO-coated microscope slides as described above, with the exception that *B. subtilis* was inoculated in ISP2 medium. One-week old sterile *Medicago sativa* roots were laid across the top of each agarose pad and incubated at RT for 14 h (for *B. subtilis*) or at 30ºC for 4 days for *S. coelicolor*.

**Sample preparation for MALDI-IMS and MALDI-MS/MS**

Samples were dried in an incubator at 65ºC for 30 minutes. A mix of calibration standards (MSCal4, Sigma-Aldrich) was spotted on each microscope slide to assess signal intensity and mass accuracy for each sample analyzed. Images were acquired using an inverted microscope (Zeiss Primovert) equipped with phase contrast objectives (100x, 200x and 400x magnification) and pictures were taken using an iPhone through the LabCam® microscope adapter from iDu Optics® (New York, NY, USA), and areas of interest framed by hand with a fine-tip lab marker. MALDI matrix was applied to the samples as described in [2] with minor modifications. Briefly, the matrix Super-DHB (Sigma-Aldrich) was sublimated under vacuum (0.05 Torr) for 5 minutes, with a temperature starting at 30ºC and ending at 95ºC, leading to matrix deposition of ~0.25-0.5 mg/cm$^2$. Samples were not rehydrated post-matrix deposition.

**MALDI-IMS and MALDI-MS/MS**

MALDI-IMS was performed in positive ion mode using a SubAP/MALDI(ng) source (MassTech, Columbia, MD). This ionization source has a small laser spot size (<10 µm) which provides high spatial resolution, and is equipped with an ion funnel, which focuses ions into the mass spectrometer, improving the signal to noise ratio [3]. This SubAP/MALDI(ng) source was coupled to a Thermo Q-Exactive (Thermo Fisher Scientific, San Jose, CA) high-resolution mass spectrometer. The full MS1 scan was done in positive mode, within a mass range of *m/z* 600-1600, resolution of 35,000 full width at half-maximum (FWHM), automatic gain control (AGC) target of 1 x $10^6$ ions and a maximum ion injection time (IT) of 400 ms. The laser energy was 50%-80% at 1 kHz repetition rate. Images were acquired using a pixel size of 10 µm at a laser velocity of 1.5 mm/min. Images were analyzed and processed using Datacube Explorer v2.3 [4], MSiReader v1.01 [5], and ImageJ v1.52a [6]. Signal intensities were adjusted to allow clear visualization.

The MS/MS spectra were acquired using the same laser parameters. The full MS1 scan was acquired within a mass range of *m/z* 600-950 for lipids and *m/z* 1000-1200 for surfactins, at a resolution of 35,000 FWHM, AGC target of 1 x $10^6$ ions and a maximum ion injection time (IT) of 1200 ms. The sequential MS/MS was performed using an inclusion list targeting the ions of interest, at a resolution of 17,500 FWHM, AGC target of 2 x $10^4$ ions, IT of 1000 ms, using an isolation window of 3 *m/z* and normalized collision energy (NCE) of 20, 30 and 45.

**Compound identification**

Compounds were identified at multiple confidence levels according to the scheme proposed by Schymanski *et al.* [7]. These results are summarized for each ID in Supplementary Table S1 and S2. Detected compounds were identified by exact mass and presence of multiple isotopes or adducts (identification level 4), allowing a maximal error of 10 ppm. When the MS2 spectra were available, the fragmentation pattern was compared to the literature (identification level 3 in the cases where diagnostic peaks made it possible to identify the lipid class) or to a commercial standard (identification level 1). A surfactin mixture was purchased from Sigma-Aldrich (S3523: Surfactin from *Bacillus subtilis*, ≥98.0%).

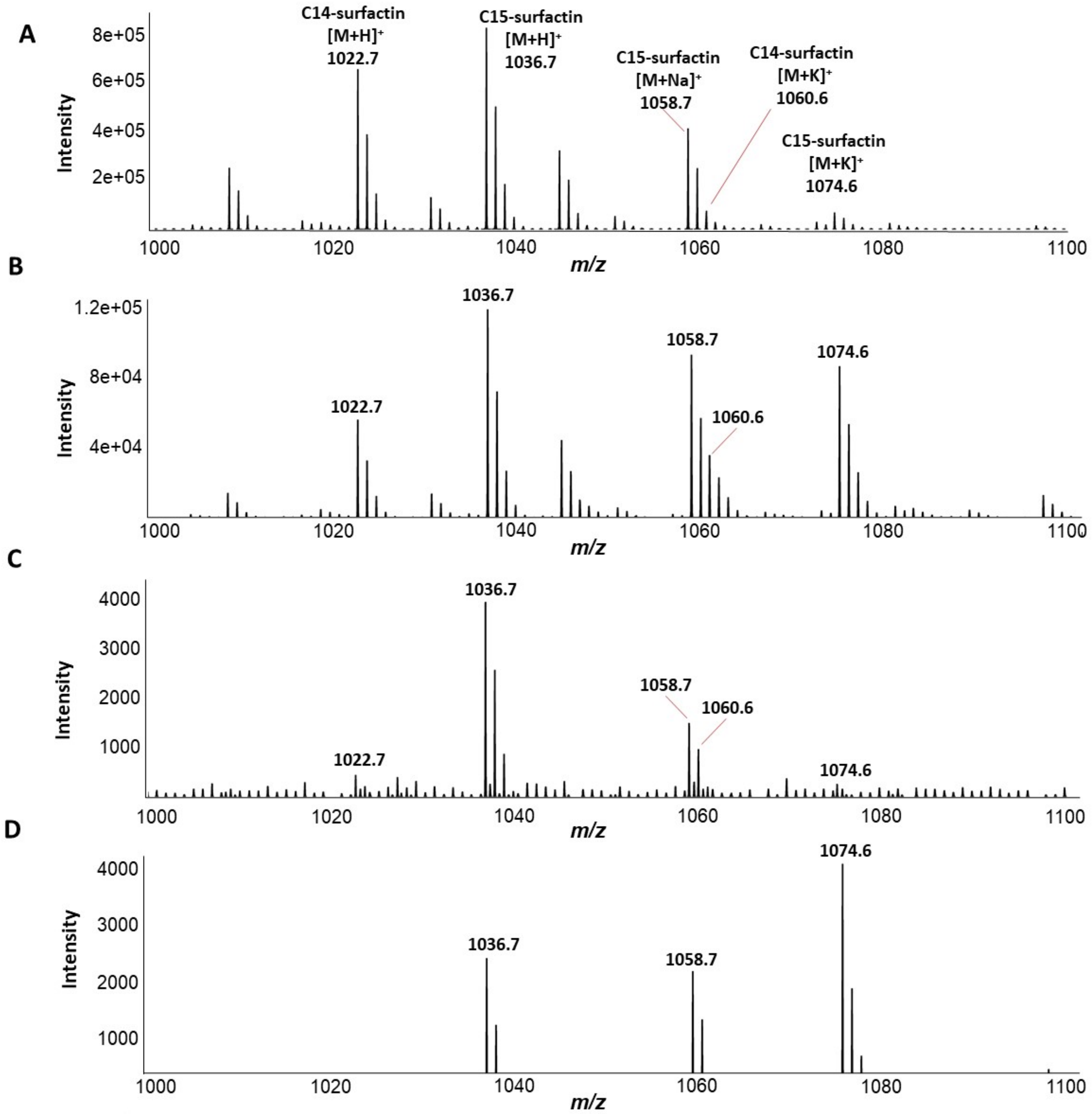
A
C14-surfactin
[M+H]+
1022.7
C15-surfactin
[M+H]+
1036.7
C15-surfactin
[M+Na]+
1058.7
C14-surfactin
[M+K]+
1060.6
C15-surfactin
[M+K]+
1074.6
Intensity
m/z
B
1036.7
1058.7
1074.6
1022.7
1060.6
C
1036.7
1058.7
1060.6
1022.7
1074.6
D
1074.6
1036.7
1058.7

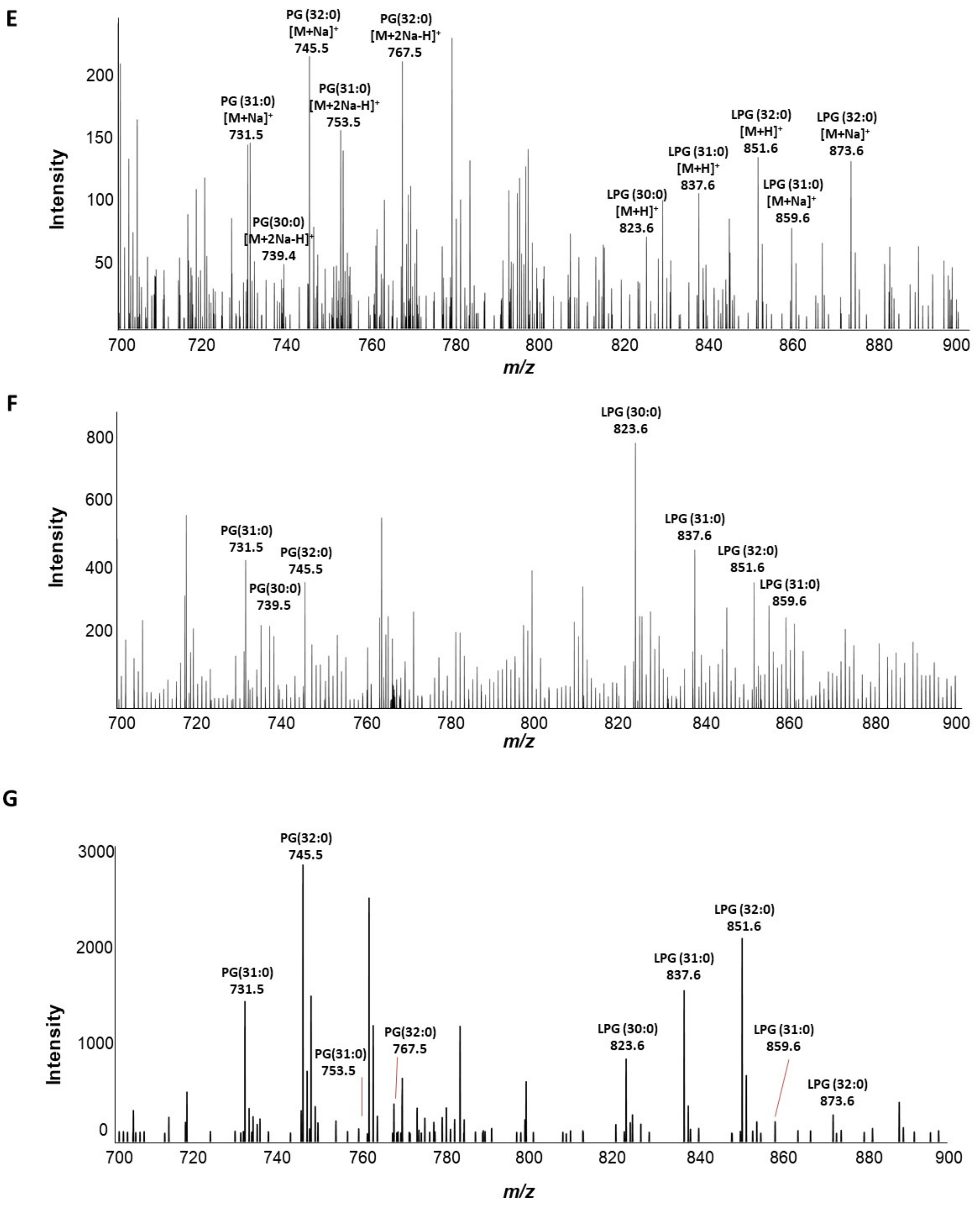


**Fig. S1:** MS1 spectra (*m/z* 1000-1100) highlighting detected surfactin analogs and adducts of a commercial surfactin standard **(A)**, a *B. subtilis* macrocolony **(B)**, a *B. subtilis* microcolony **(C)** and *B. subtilis* microcolonies next to a *M. sativa* root **(D)**. MS1 spectra (*m/z* 700-900) highlighting detected lipids of a lawn of *B. subtilis* microcolonies **(E)**, *B. subtilis* microcolony **(F)** and *B. subtilis* microcolonies next to *M. sativa* root **(G)**.

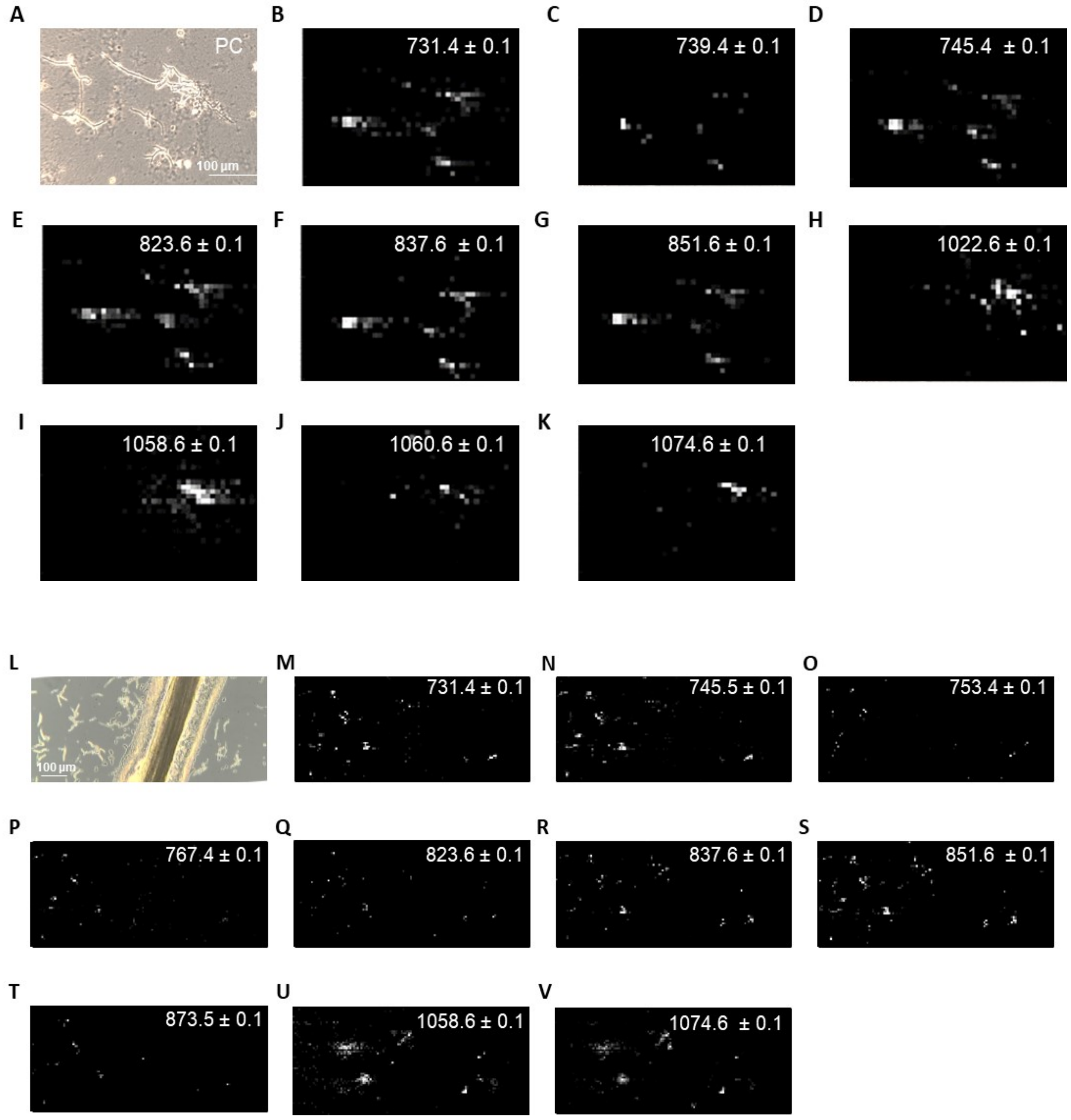


**Fig. S2:** Micrograph of *B. subtilis* microcolonies **(A)** and the corresponding MALDI images showing the spatial distribution of identified lipids **(B-G)** and surfactins **(H-K)**. Scale bar = 100 microns. Micrograph of *B. subtilis* microcolonies next to *M. sativa* root **(L)** and the corresponding MALDI images showing the spatial distribution of identified lipids **(M-T)** and surfactins **(U,V)**. Scale bar = 100 microns. All micrographs were acquired using a phase contrast (PC) microscope.

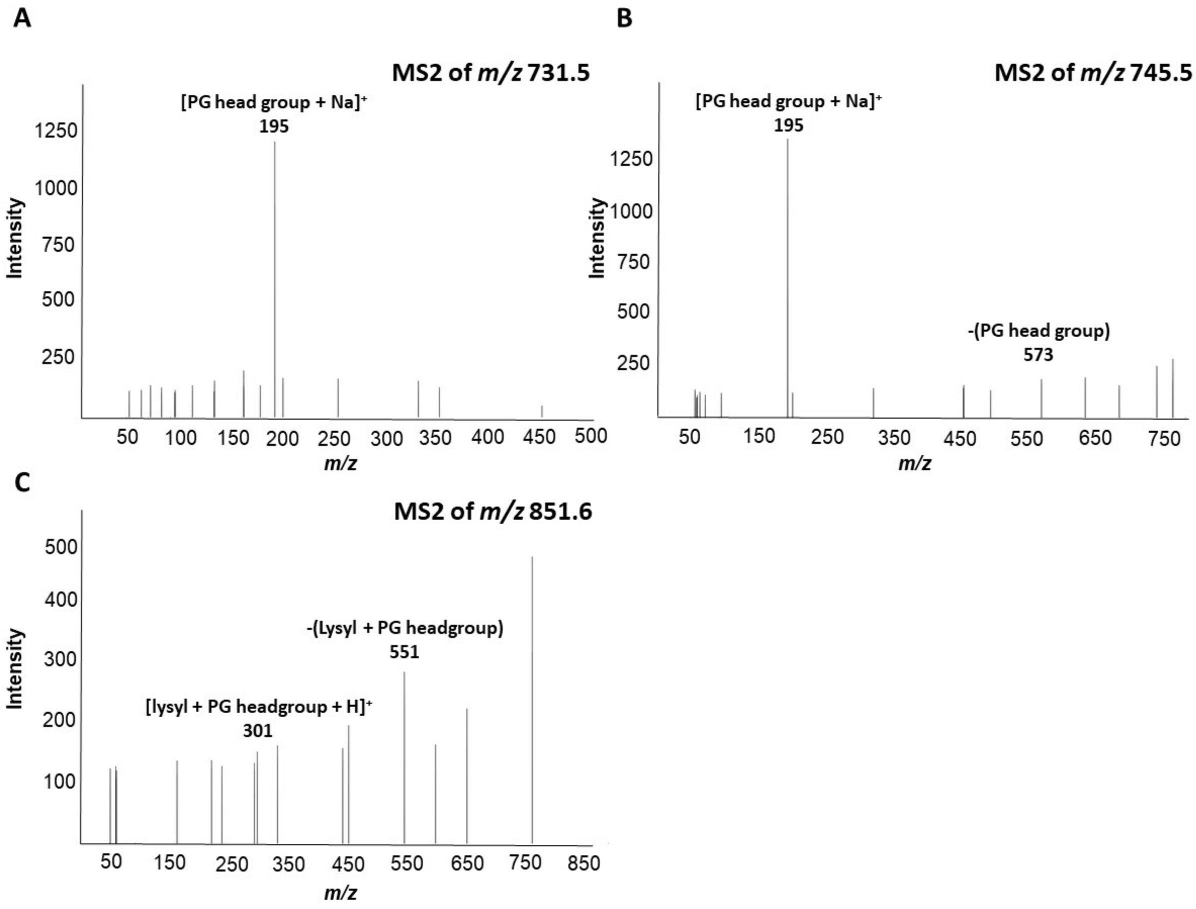


**Fig. S3:** MS2 spectra of level 3 identified *B. subtilis* PG and LPG lipids. PG (31:0) $[M+Na]^+$ **(A)**. PG (32:0) $[M+Na]^+$ **(B)**. LPG (32:0) $[M+H]^+$ **(C)**. All MS2 spectra were acquired using NCE of 20.

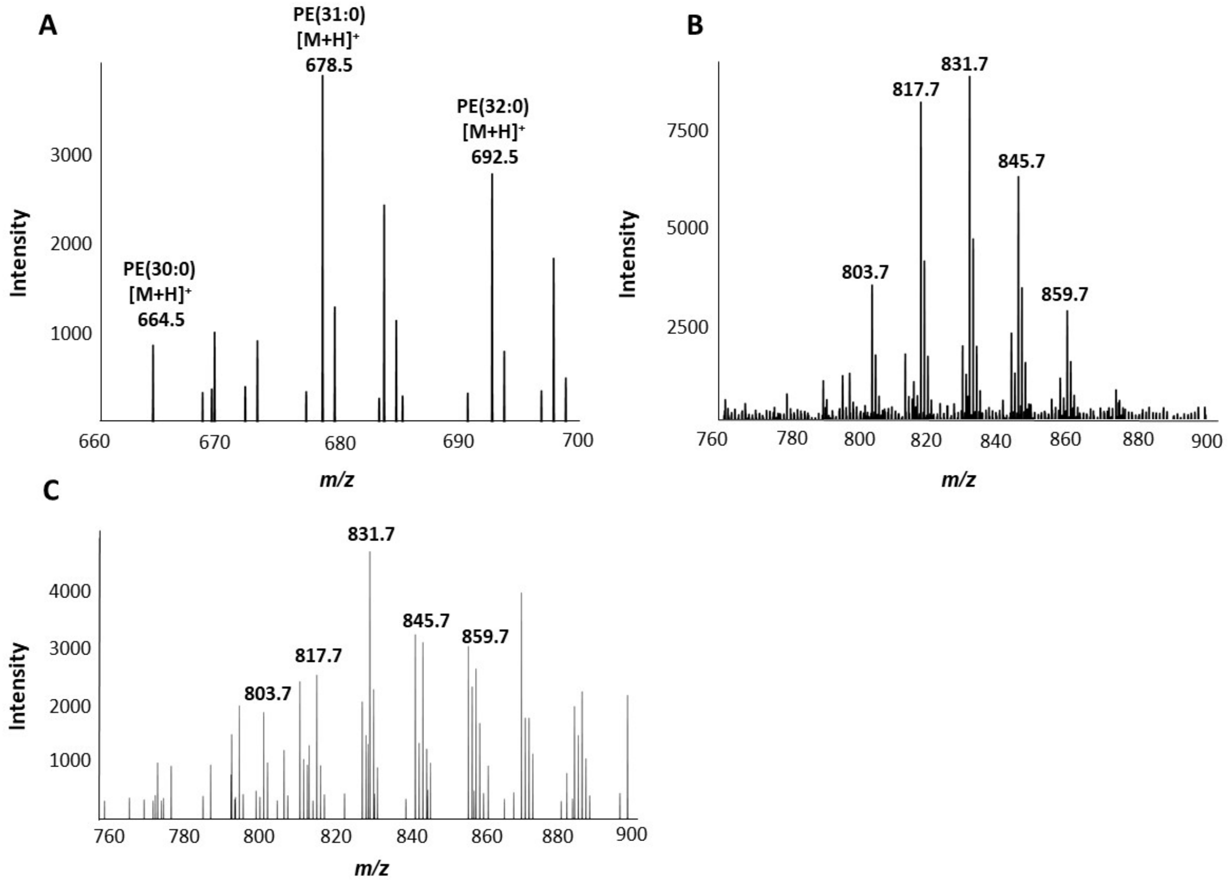


**Fig. S4:** MS1 spectra (*m/z* 660-700) of *S. coelicolor* microcolonies highlighting level 4 identified PE lipids **(A)**. MS1 spectra (*m/z* 760-900) of *S. coelicolor* microcolonies highlighting uncharacterized lipids from a *S. coelicolor* microcolony **(B)** and a *S. coelicolor* microcolony next to a *M. sativa* root **(C)**.

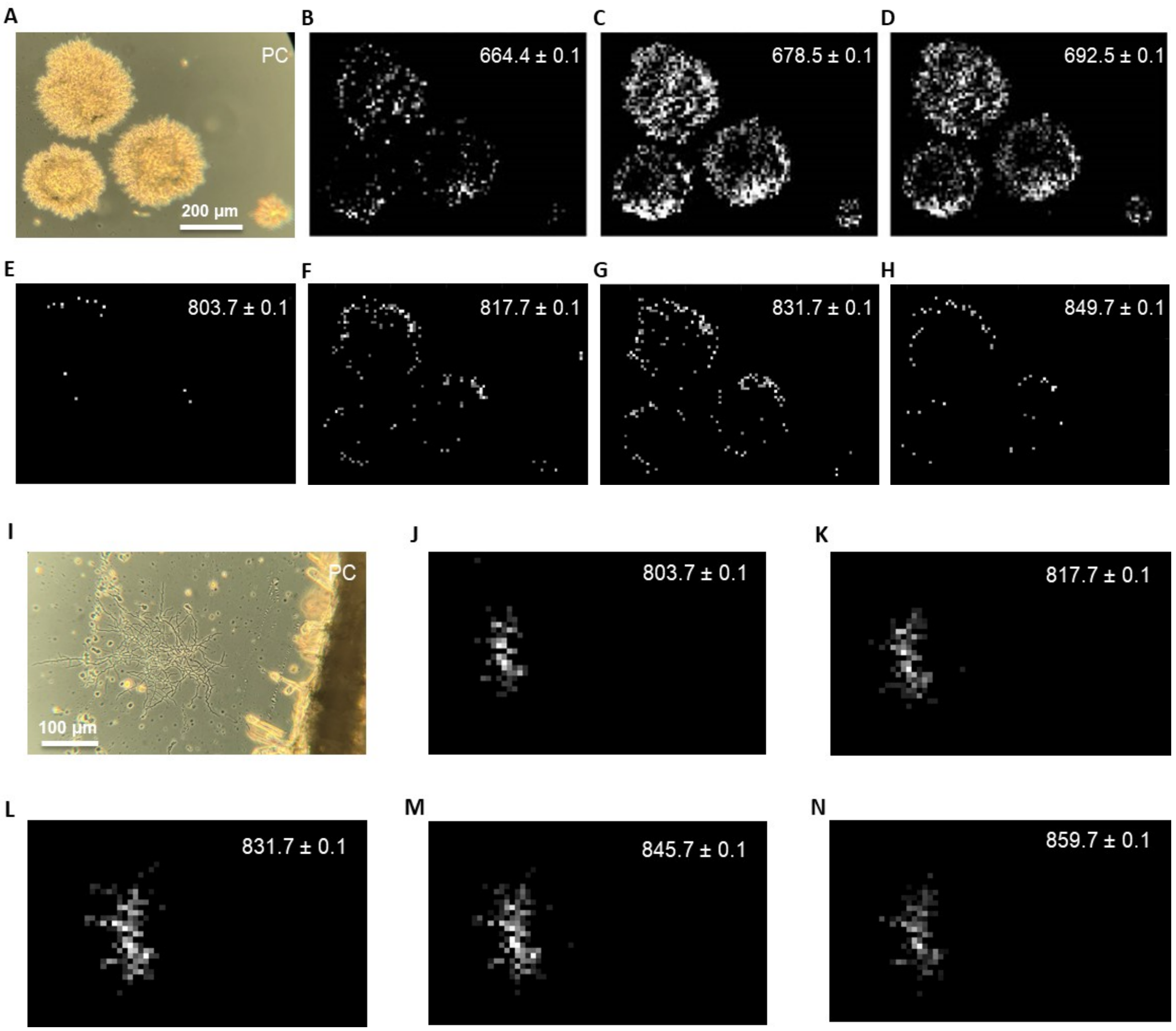


**Fig. S5:** Micrograph of *S. coelicolor* microcolonies **(A)** and the corresponding MALDI images showing the spatial distribution of level 4 identified PE lipids **(B-D)** and uncharacterized lipids **(E-H).** Scale bar = 200 microns. Micrograph of *S. coelicolor* microcolonies next to *M. sativa* root **(I)** and the corresponding MALDI images showing the spatial distribution of uncharacterized lipids **(J-N)**. Scale bar = 100 microns. All micrographs were acquired using a phase contrast (PC) microscope.

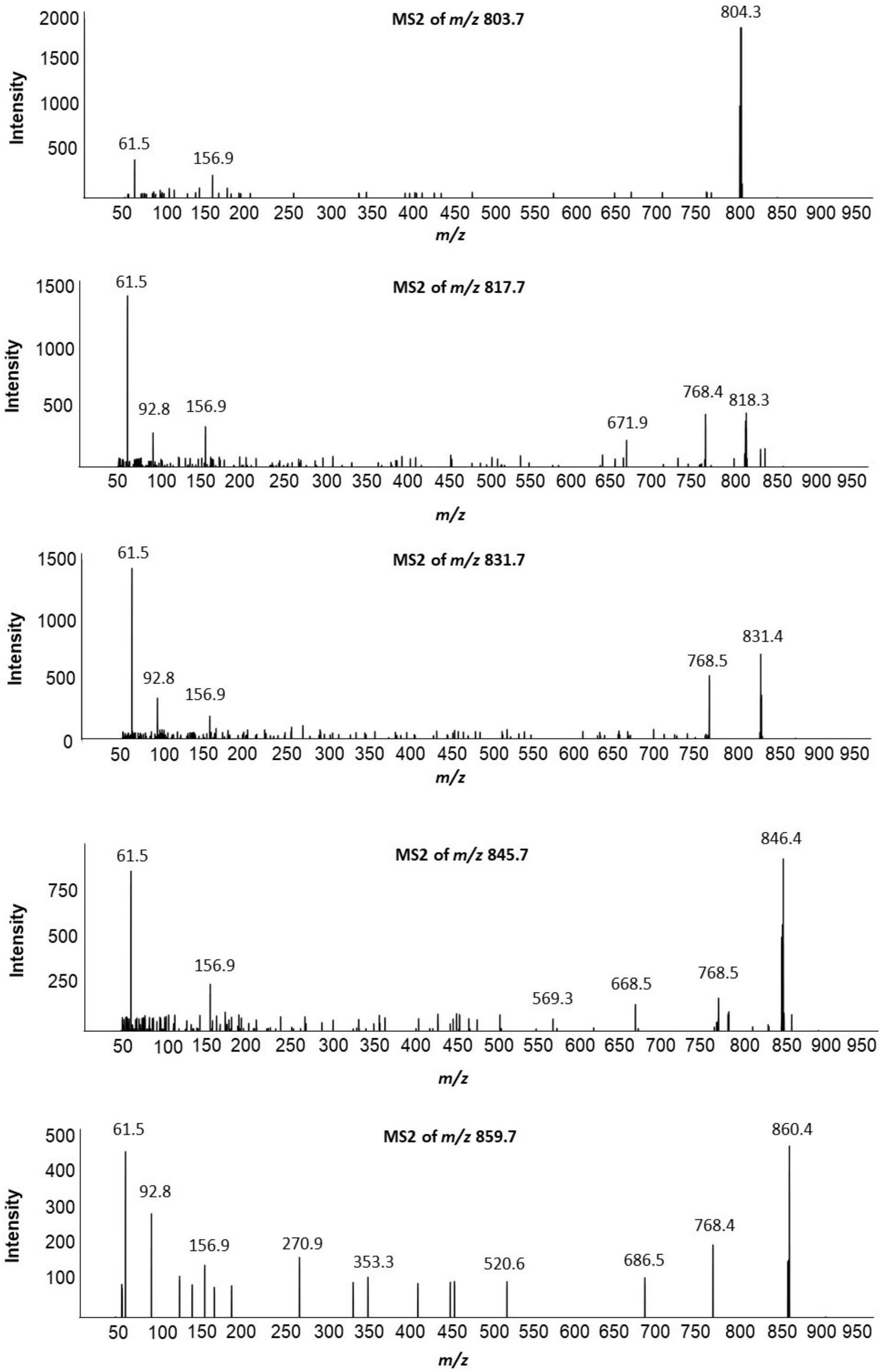


**Fig. S6:** MS2 spectra of uncharacterized *S. coelicolor* lipids. All MS2 spectra were acquired using NCE of 30.

**Table S1:** Identification of features detected on *B. subtilis* and/or *B. subtilis* + *M. sativa* root MALDI-IMS experiments, showing the putative ID, calculated ppm error, how each feature was identified, and the respective MALDI-IMS that was acquired. Gidden *et al.*, Samant *et al.*, Aleti *et al.*, and Debois *et al.* are references [8], [9], [10], and [11], respectively.

| | | | | | **Identification basis** | | | | | |
|---|---|---|---|---|---|---|---|---|---|---|
| **Putative ID** | **Source** | **Observed *m/z*** | **Calculated *m/z*** | **ppm error** | **MS1** | **MS2** | **Literature** | **Commercial Standard** | **Identification level** | **MALDI Image** |
| PG (31:0)<br>$[M+Na]^+$ | BS-LMi<br>BS-Mi<br>BS+R | 731.484563<br>731.481991<br>731.487127 | 731.483356 | 1.7<br>1.9<br>5.2 | Fig. S1E<br>Fig. S1F<br>Fig. S1G | Fig. S3A<br>NA<br>NA | Gidden *et al.*, 2009 | -- | 3 | NA<br>Fig. S2B<br>Fig. S2M |
| PG (30:0)<br>$[M+2Na-H]^+$ | BS-LMi<br>BS-Mi<br>BS+R | 739.448992<br>739.455475<br>ND | 739.449648 | 0.9<br>7.9<br>-- | Fig. S1E<br>Fig. S1F<br>-- | ND<br>NA<br>NA | Gidden *et al.*, 2009 | -- | 4 | NA<br>Fig. S2C<br>-- |
| PG (32:0)<br>$[M+Na]^+$ | BS-LMi<br>BS-Mi<br>BS+R | 745.502587<br>745.499737<br>745.495619 | 745.499006 | 4.8<br>1.0<br>4.5 | Fig. S1E<br>Fig. S1F<br>Fig. S1G | Fig. S3B<br>NA<br>NA | Gidden *et al.*, 2009 | -- | 3 | NA<br>Fig. S2D<br>Fig. S2N |
| PG (31:0)<br>$[M+2Na-H]^+$ | BS-LMi<br>BS-Mi<br>BS+R | 753.462233<br>ND<br>753.464664 | 753.465298 | 4.1<br>--<br>0.8 | Fig. S1E<br>--<br>Fig. S1G | ND<br>NA<br>NA | Gidden *et al.*, 2009 | -- | 4 | NA<br>--<br>Fig. S2O |
| PG (32:0)<br>$[M+2Na-H]^+$ | BS-LMi<br>BS-Mi<br>BS+R | 767.483854<br>ND<br>767.476709 | 767.480948 | 3.8<br>--<br>5.5 | Fig. S1E<br>--<br>Fig. S1G | ND<br>--<br>NA | Gidden *et al.*, 2009 | -- | 4 | NA<br>--<br>Fig. S2P |
| LPG (30:0)<br>$[M+H]^+$ | BS-LMi<br>BS-Mi<br>BS+R | 823.580263<br>823.576782<br>823.58469 | 823.580727 | 0.6<br>4.8<br>4.8 | Fig. S1E<br>Fig. S1F<br>Fig. S1G | ND<br>NA<br>NA | Gidden *et al.*, 2009 | -- | 4 | NA<br>Fig. S2E<br>Fig. S2Q |
| LPG (31:0)<br>$[M+H]^+$ | BS-LMi<br>BS-Mi<br>BS+R | 837.598111<br>837.595454<br>837.592319 | 837.596377 | 2.1<br>1.1<br>4.8 | Fig. S1E<br>Fig. S1F<br>Fig. S1G | ND<br>NA<br>NA | Samant *et al.*, 2009 | -- | 4 | NA<br>Fig. S2F<br>Fig. S2R |
| LPG (32:0)<br>$[M+H]^+$ | BS-LMi<br>BS-Mi<br>BS+R | 851.612318<br>851.609664<br>851.606701 | 851.612027 | 0.3<br>2.8<br>6.3 | Fig. S1E<br>Fig. S1F<br>Fig. S1G | Fig. S3C<br>NA<br>NA | Gidden *et al.*, 2009 | -- | 3 | NA<br>Fig. S2G<br>Fig. S2S |
| LPG (31:0)<br>$[M+Na]^+$ | BS-LMi<br>BS-Mi<br>BS+R | 859.577326<br>859.573062<br>859.582151 | 859.578319 | 1.2<br>6.1<br>4.5 | Fig. S1E<br>Fig. S1F<br>Fig. S1G | ND<br>NA<br>NA | -- | -- | 4 | NA<br>NS<br>NS |
| LPG(32:0)<br>$[M+Na]^+$ | BS-LMi<br>BS-Mi<br>BS+R | 873.598345<br>ND<br>873.590877 | 873.593969 | 5.0<br>--<br>3.5 | Fig. S1E<br>--<br>Fig. S1G | ND<br>NA<br>NA | -- | -- | 4 | NA<br>--<br>Fig. S2T |
| C14-surfactin<br>$[M+H]^+$ | BS-Ma<br>BS-Mi<br>BS+R<br>S-Std | 1022.677755<br>1022.677504<br>ND<br>1022.676221 | 1022.674764 | 2.9<br>2.7<br>--<br>1.4 | Fig. S1B<br>Fig. S1C<br>--<br>Fig. S1A | NS<br>NA<br>NA<br>NS | Aleti *et al.*, 2016 | Surfactin (S3523,Sigma-Aldrich) | 1 | NA<br>Fig. S2H<br>--<br>NA |
| C15-surfactin | BS-Ma<br>BS-Mi | 1036.685225<br>1036.68503 | 1036.690413 | 5.0<br>5.2 | Fig. S1B<br>Fig. S1C | Fig. 1B<br>NA | Aleti *et al.*, 2016 | Surfactin (S3523,Sigma- | 1 | NA<br>Fig. 1A |

| | | | | | | | | | | |
|---|---|---|---|---|---|---|---|---|---|---|
| [M+H]$^+$ | BS+R | 1036.682429 | | 7.7 | Fig. S1D | NA | | Aldrich) | | Fig. 2A |
| | S-Std | 1036.684068 | | 6.1 | Fig. S1A | Fig 1B | | | | NA |
| C15-surfactin [M+Na]$^+$ | BS-Ma | 1058.666837 | 1058.672355 | 5.2 | Fig. S1B | -- | Debois *et al.*, 2014 | Surfactin (S3523,Sigma-Aldrich) | 1 | NA |
| | BS-Mi | 1058.666731 | | 5.3 | Fig. S1C | NA | | | | Fig. S2I |
| | BS+R | 1058.664453 | | 7.5 | Fig. S1D | NA | | | | Fig. S2U |
| | S-Std | 1058.664917 | | 7.0 | Fig. S1A | NS | | | | NA |
| C14-surfactin [M+K]$^+$ | BS-Ma | 1060.629763 | 1060.630646 | 0.8 | Fig. S1B | NS | Debois *et al.*, 2014 | Surfactin (S3523,Sigma-Aldrich) | 1 | NA |
| | BS-Mi | 1060.629666 | | 0.9 | Fig. S1C | NA | | | | Fig. S2J |
| | BS+R | ND | | -- | -- | -- | | | | -- |
| | S-Std | 1060.627841 | | 2.6 | Fig. S1A | NS | | | | NA |
| C15-surfactin [M+K]$^+$ | BS-Ma | 1074.641328 | 1074.646295 | 4.6 | Fig. S1B | NS | Debois *et al.*, 2014 | Surfactin (S3523,Sigma-Aldrich) | 1 | NA |
| | BS-Mi | 1074.641289 | | 4.7 | Fig. S1C | NA | | | | Fig. S2K |
| | BS+R | 1074.639253 | | 6.6 | Fig. S1D | NA | | | | Fig. S2V |
| | S-Std | 1074.652469 | | 5.7 | Fig. S1A | NS | | | | NA |

ND = not detected

NA = not acquired

NS = not shown

-- = not applicable

PG = Phosphatidylglycerol

LPG = Lysyl-phosphatidylglycerol

BS-LMi = *B. subtilis* lawn of microcolonies

BS-Ma = *B. subtilis* macrocolony

BS-Mi = *B. subtilis* microcolony

BS+R = *B. subtilis* microcolony next to a *M. sativa* root

S-Std = Surfactin standard

**Table S2** – Identification of features detected on *S. coelicolor* microcolonies MALDI-IMS experiment, showing the putative ID, calculated ppm error, how each feature was identified, and the respective MALDI-IMS that was acquired. Sandoval-Calderón is reference [12].

| | | | | | Identification basis | | | | |
|---|---|---|---|---|---|---|---|---|---|
| **Putative ID** | **Source** | **Observed *m/z*** | **Calculated *m/z*** | **ppm error** | **MS1** | **MS2** | **Literature** | **Identification level** | **MALDI image** |
| PE(30:0) $[M+H]^+$ | Sco-Mi | 664.488877 | 664.491183 | 3.5 | Fig. 4A | ND | Sandoval-Calderón *et al.* 2015 | 4 | Fig. S5B |
| PE(31:0) $[M+H]^+$ | Sco-Mi | 678.50798 | 678.506833 | 1.7 | Fig. 4A | ND | Sandoval-Calderón *et al.* 2015 | 4 | Fig. S5C |
| PE(32:0) $[M+H]^+$ | Sco-Mi | 692.524058 | 692.522481 | 2.3 | Fig. 4A | ND | Sandoval-Calderón *et al.* 2015 | 4 | Fig. S5D |

Sco-Mi = *S. coelicolor* microcolony
PE = Phosphatidylethanolamine
